# Angiomotin isoform 2 promotes binding of PALS1 to KIF13B at the base of primary cilia and suppresses ciliary elongation

**DOI:** 10.1101/2021.10.14.464392

**Authors:** Stine Kjær Morthorst, Camilla Nielsen, Pietro Farinelli, Zeinab Anvarian, Christina Birgitte R. Rasmussen, Andrea Serra-Marques, Ilya Grigoriev, Maarten Altelaar, Nicoline Fürstenberg, Alexander Ludwig, Anna Akhmanova, Søren Tvorup Christensen, Lotte Bang Pedersen

## Abstract

The kinesin-3 motor KIF13B functions in endocytosis, vesicle transport, and regulation of ciliary length and signaling. Direct binding of the membrane-associated guanylate kinase (MAGUK) DLG1 to KIF13B’s MAGUK-binding stalk (MBS) domain relieves motor autoinhibition and promotes microtubule plus end-directed cargo transport. Here we characterize Angiomotin isoform 2 (Ap80) as a novel KIF13B interactor that promotes binding of another MAGUK, the polarity protein and Crumbs complex component PALS1, to KIF13B. Live cell imaging analysis indicated that Ap80 is concentrated at the base of primary cilia and recruits PALS1 to this site, but is not itself a cargo of KIF13B. Consistent with a ciliary function for Ap80, its depletion led to elongated primary cilia while its overexpression caused ciliary shortening. Our results suggest that Ap80 may specifically activate KIF13B cargo binding at the base of primary cilia to regulate ciliary length.

## Introduction

Kinesin motors mediate microtubule-based transport of diverse intracellular cargoes, including organelles and vesicles (Hirokawa and Tanaka, 2015). In humans 8 kinesin genes code for kinesin-3 family members, which are highly processive microtubule plus-end directed motors involved in vesicle transport and endocytosis (Siddiqui and Straube, 2017). Kinesin-3 member KIF13B (also known as GAKIN; guanylate kinase-associated kinesin) was initially identified as a direct binding partner of the scaffold protein DLG1 in lymphocytes (Hanada et al., 2000). Subsequent studies implicated KIF13B in multiple cellular and physiological processes e.g. regulation of neuronal polarity and axon formation (Horiguchi et al., 2006, Yoshimura et al., 2010); Schwann cell and oligodendrocyte myelination (Bolis et al., 2009, Noseda et al., 2016); cholesterol uptake and CAV1-dependent endocytosis of LRP1 in hepatocytes (Kanai et al., 2014, Mills et al., 2019); Golgi to plasma membrane trafficking of VEGFR2 in endothelial cells (Yamada et al., 2014) and of Rab6 exocytotic vesicles in HeLa cells (Serra-Marques et al., 2020); germ cell migration and Planar Cell Polarity signaling in *Xenopus laevis* (Tarbashevich et al., 2011, Ossipova et al., 2015); and regulation of ciliary CAV1 distribution and Sonic hedgehog (Shh) signaling in hTERT-immortalized retinal pigment epithelial (hereafter RPE1) cells (Schou et al., 2017).

KIF13B function requires tight temporal and spatial control of its motor activity, which is mediated by binding partners that recognize specific domains in KIF13B to regulate its conformation and dimerization (Soppina et al., 2014, Siddiqui and Straube, 2017, Morthorst et al., 2018, Ren et al., 2018). The best characterized KIF13B regulator and cargo is DLG1, a Scribble polarity complex component of the membrane-associated guanylate kinase (MAGUK) homologue family. DLG1 binds directly to the MAGUK-binding stalk (MBS) domain of KIF13B, which relieves motor autoinhibition and promotes KIF13B-mediated transport of DLG1 along microtubules (Hanada et al., 2000, Asaba et al., 2003, Yamada et al., 2007, Zhu et al., 2016). Similarly, other MAGUKs including DLG4 and PALS1, a core component of the Crumbs complex required for development and maintenance of epithelial cell polarity in diverse tissues, bind directly to the KIF13B MBS domain (Zhu et al., 2016). Furthermore, we identified an interaction between KIF13B and the ciliary transition zone (TZ) protein NPHP4 (Schou et al., 2017), which also associates with PALS1 and PATJ (Sang et al., 2011, Delous et al., 2009). However, it is unknown whether interaction between KIF13B and PALS1 is relevant in a ciliary context.

Primary cilia are microtubule-based sensory organelles present on the surface of many vertebrate cell types, where they coordinate e.g. Shh (Bangs and Anderson, 2017) and receptor tyrosine kinase signaling (Christensen et al., 2017). The ciliary axoneme extends from the basal body and is surrounded by a membrane, which is continuous with but compositionally distinct from the plasma membrane. This compartmentalization is crucial for ciliary function, and is regulated by the TZ at the ciliary base (Garcia-Gonzalo and Reiter, 2017), as well as by intraflagellar transport (IFT) that via kinesin-2 and dynein 2 motors, respectively, moves trains of IFT particles with ciliary cargoes up and down the cilum (Taschner and Lorentzen, 2016). Nematodes additionally express a kinesin-3 motor, KLP-6, that regulates ciliary membrane composition and function specifically in male sensory neurons by promoting release of extracellular vesicles from the ciliary tip (Akella and Barr, 2021). Recently, we demonstrated that KIF13B displays bursts of bidirectional intraciliary movement within primary cilia of RPE1 cells and is occasionally released from the ciliary tip (Juhl et al., 2021). How KIF13B intraciliary movement is regulated and the identity of ciliary cargoes of KIF13B are not known.

Here we characterize Angiomotin isoform 2 (Ap80) as a KIF13B interactor and show that Ap80 promotes the binding of KIF13B to PALS1. We show that Ap80 is concentrated at and recruits PALS1 to the ciliary base of RPE1 cells, and that its depletion and overexpression cause ciliary elongation and shortening, respectively. Thus Ap80 may activate KIF13B cargo binding at the ciliary base to regulate ciliary length.

## Results and discussion

### KIF13B interacts with the N-terminus of Ap80

To identify KIF13B interactors we performed streptavidin pull-down assays with BirA- and GFP-tagged (BirAGFP) KIF13B constructs combined with mass spectrometry (Table 1). We used the deletion mutants BirAGFP-KIF13B Tail 2 (residues 607-1826) and BirAGFP-KIF13B Tail 3 (residues 752-1826), which contain and lack the MBS domain, respectively (Serra-Marques et al., 2020), reasoning that these may reveal MBS-specific interactors. We identified several significant hits that had more peptides in the Tail 2 mutant sample compared to Tail 3 (Table 1). These included DLG1, and its known interactors MPP7, LIN7C and CASK (Bohl et al., 2007, Lee et al., 2002), and utrophin, a large cytoskeletal adaptor that binds to KIF13B in a complex mediating endocytosis of LRP1 (Kanai et al., 2014). Known interactors of utrophin, syntrophin (Kramarcy et al., 1994) and dystrobrevins (Peters et al., 1997), were also present in the pull down. The most prominent potential novel binding partners of KIF13B, showing significantly stronger association with Tail 2 compared to Tail 3, were kinase D-interacting substrate of 220 kDa (KIDINS220), a conserved transmembrane molecule implicated in signaling (Neubrand et al., 2012), and an adaptor protein belonging to the motin family, angiomotin (AMOT) (Moleirinho et al., 2014) (Table 1).

**Table 1.**
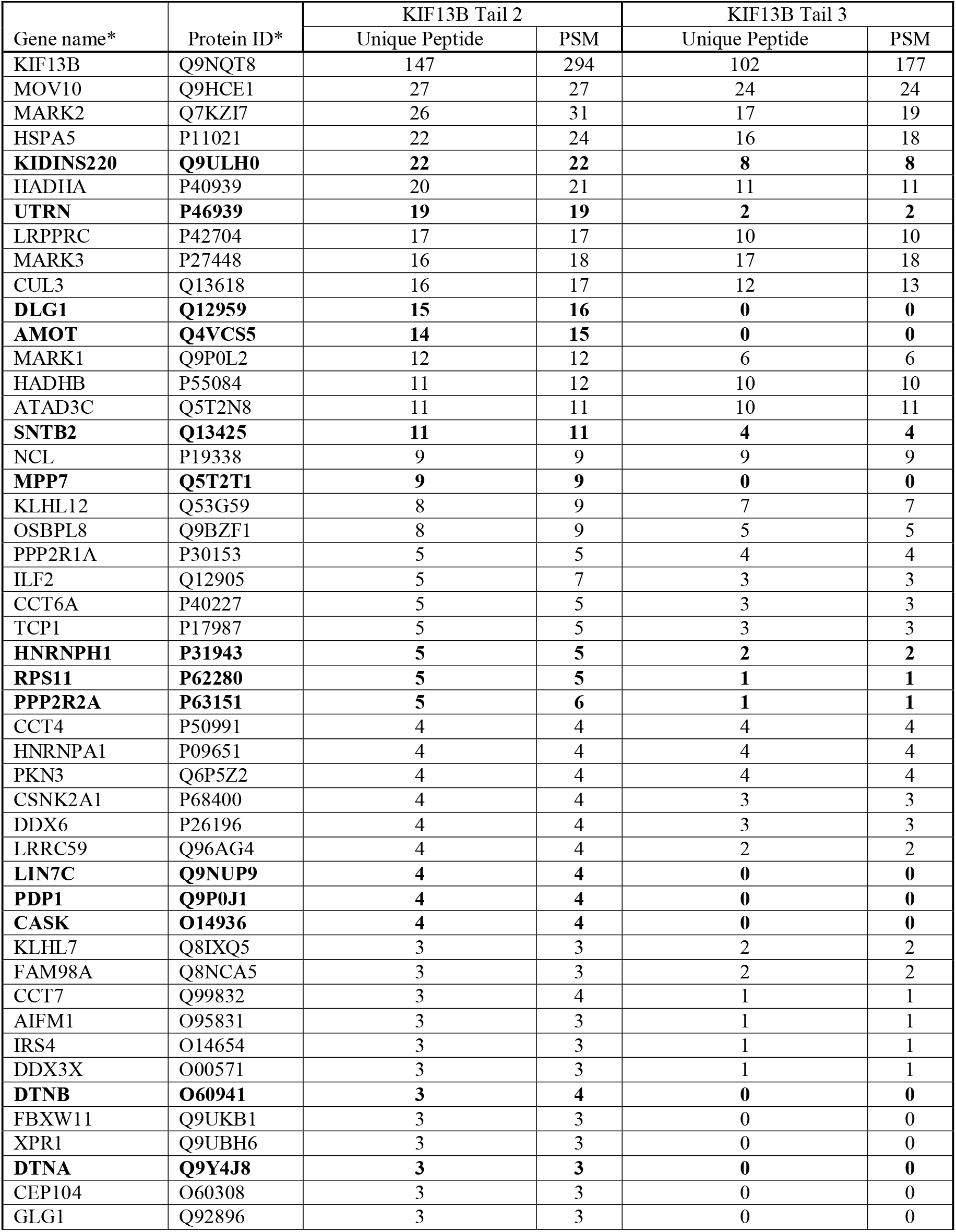
Binding partners of the indicated BirAGFP-KIF13B constructs in HEK293T cells identified by mass spectrometry analysis. ^*^The highlighted genes/proteins preferentially bind the KIF13B Tail 2 over Tail 3 and are discussed in the text. Only proteins identified by at least three unique peptides in the BirAGFP-KIF13B Tail 2 sample are included. PSM, peptide spectrum matches.

| Gene name* | Protein ID* | KIF13B Tail 2 |  | KIF13B Tail 3 |  |
| --- | --- | --- | --- | --- | --- |
|  |  | Unique Peptide | PSM | Unique Peptide | PSM |
| KIF13B | Q9NQT8 | 147 | 294 | 102 | 177 |
| MOV10 | Q9HCE1 | 27 | 27 | 24 | 24 |
| MARK2 | Q7KZI7 | 26 | 31 | 17 | 19 |
| HSPA5 | P11021 | 22 | 24 | 16 | 18 |
| **KIDINS220** | **Q9ULH0** | **22** | **22** | **8** | **8** |
| HADHA | P40939 | 20 | 21 | 11 | 11 |
| **UTRN** | **P46939** | **19** | **19** | **2** | **2** |
| LRPPRC | P42704 | 17 | 17 | 10 | 10 |
| MARK3 | P27448 | 16 | 18 | 17 | 18 |
| CUL3 | Q13618 | 16 | 17 | 12 | 13 |
| **DLG1** | **Q12959** | **15** | **16** | **0** | **0** |
| **AMOT** | **Q4VCS5** | **14** | **15** | **0** | **0** |
| MARK1 | Q9P0L2 | 12 | 12 | 6 | 6 |
| HADHB | P55084 | 11 | 12 | 10 | 10 |
| ATAD3C | Q5T2N8 | 11 | 11 | 10 | 11 |
| **SNTB2** | **Q13425** | **11** | **11** | **4** | **4** |
| NCL | P19338 | 9 | 9 | 9 | 9 |
| **MPP7** | **Q5T2T1** | **9** | **9** | **0** | **0** |
| KLHL12 | Q53G59 | 8 | 9 | 7 | 7 |
| OSBPL8 | Q9BZF1 | 8 | 9 | 5 | 5 |
| PPP2R1A | P30153 | 5 | 5 | 4 | 4 |
| ILF2 | Q12905 | 5 | 7 | 3 | 3 |
| CCT6A | P40227 | 5 | 5 | 3 | 3 |
| TCP1 | P17987 | 5 | 5 | 3 | 3 |
| **HNRNPH1** | **P31943** | **5** | **5** | **2** | **2** |
| **RPS11** | **P62280** | **5** | **5** | **1** | **1** |
| **PPP2R2A** | **P63151** | **5** | **6** | **1** | **1** |
| CCT4 | P50991 | 4 | 4 | 4 | 4 |
| HNRNPA1 | P09651 | 4 | 4 | 4 | 4 |
| PKN3 | Q6P5Z2 | 4 | 4 | 4 | 4 |
| CSNK2A1 | P68400 | 4 | 4 | 3 | 3 |
| DDX6 | P26196 | 4 | 4 | 3 | 3 |
| LRRC59 | Q96AG4 | 4 | 4 | 2 | 2 |
| **LIN7C** | **Q9NUP9** | **4** | **4** | **0** | **0** |
| **PDP1** | **Q9P0J1** | **4** | **4** | **0** | **0** |
| **CASK** | **O14936** | **4** | **4** | **0** | **0** |
| KLHL7 | Q8IXQ5 | 3 | 3 | 2 | 2 |
| FAM98A | Q8NCA5 | 3 | 3 | 2 | 2 |
| CCT7 | Q99832 | 3 | 4 | 1 | 1 |
| AIFM1 | O95831 | 3 | 3 | 1 | 1 |
| IRS4 | O14654 | 3 | 3 | 1 | 1 |
| DDX3X | O00571 | 3 | 3 | 1 | 1 |
| **DTNB** | **O60941** | **3** | **4** | **0** | **0** |
| FBXW11 | Q9UKB1 | 3 | 3 | 0 | 0 |
| XPR1 | Q9UBH6 | 3 | 3 | 0 | 0 |
| **DTNA** | **Q9Y4J8** | **3** | **3** | **0** | **0** |
| CEP104 | O60308 | 3 | 3 | 0 | 0 |
| GLG1 | Q92896 | 3 | 3 | 0 | 0 |

We focused on AMOT due to its reported interaction with PALS1 (Wells et al., 2006, Tan et al., 2020), which associates with KIF13B (Zhu et al., 2016) and the KIF13B interactor NPHP4 (Delous et al., 2009, Schou et al., 2017). AMOT is found in two splice variants; angiomotin isoform 1 (Ap130) contains a 409 amino acid N-terminal extension absent in isoform 2, Ap80 (Ernkvist et al., 2006). We previously reported that KIF13B binds Ap80 (Schou et al., 2017), but the interaction was not characterized in detail. Here, we confirmed by FLAG immunoprecipitation (IP) in HEK293T cells that FLAG-Ap80 interacts with HA-tagged full-length or motorless KIF13B (KIF13B-Δmotor-HA; residues 393-1826), but FLAG-Ap130 does not (Figure 1A). Supportively, FLAG-Ap80 co-IPed with GFP-KIF13B (Figure 1B). IP with different FLAG-Ap80 truncations identified the first 34 residues in Ap80 as essential for KIF13B binding, since Ap80-NB (residues 1-245) comprising the N-terminus and BAR domain region, bound to KIF13B-Δmotor-HA, but Ap80-B (residues 35-245) did not (Figure 1C, D). Ap80 residues 1-245 are also present in Ap130, which failed to bind KIF13B (Figure 1A, C), suggesting that the KIF13B binding site in AMOT is blocked by the extended N-terminus of Ap130.

**Figure 1.**
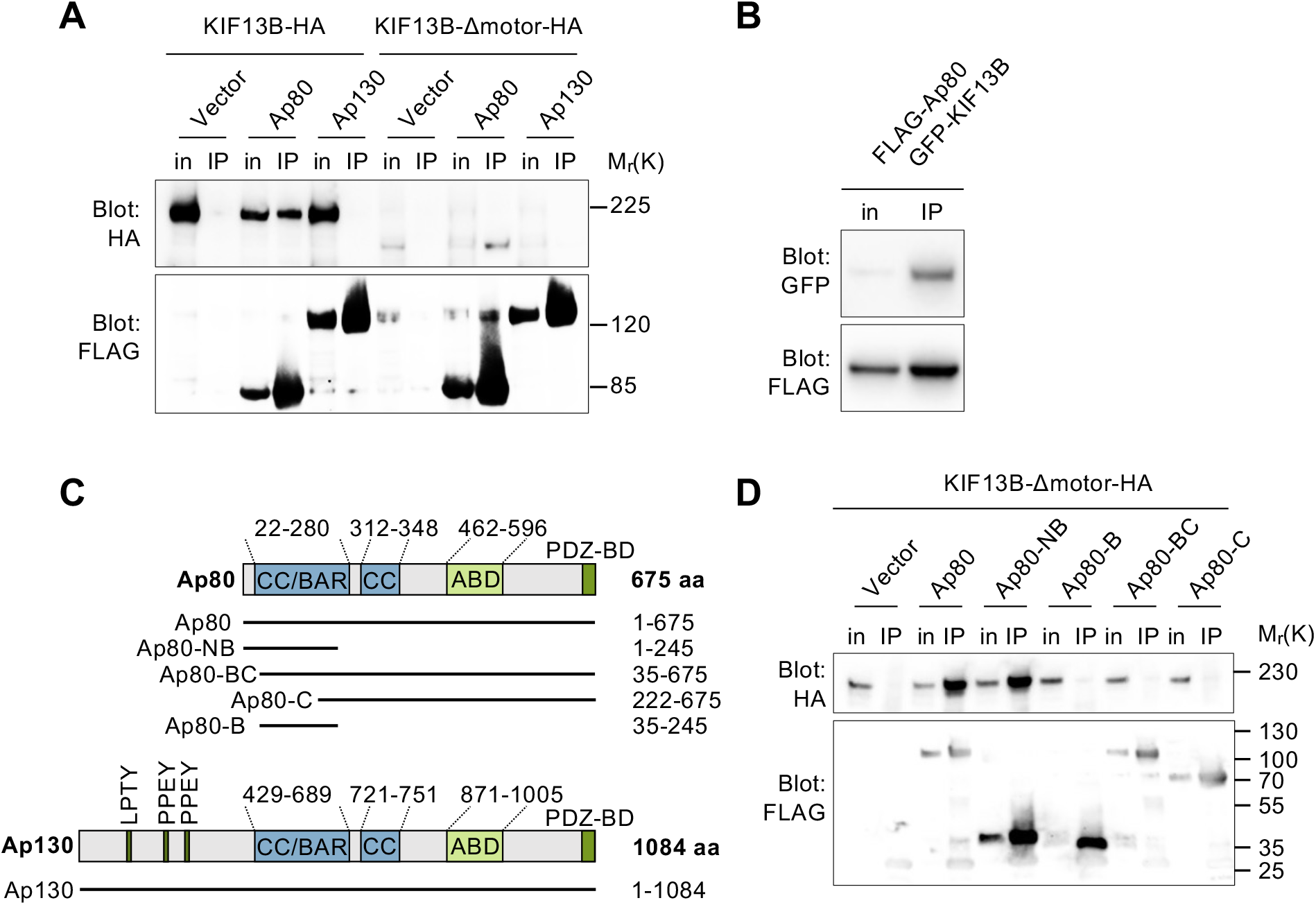
Ap80 interacts with KIF13B via its N-terminus. (**A, D**) FLAG IP of cells co-expressing FLAG-tagged Ap80 or Ap130 fusions and KIF13B-HA or KIF13B-Δmotor-HA. (**B)** GFP IP of cells co-expressing FLAG-Ap80 and GFP-KIF13B. Input (in) and IP pellet (IP) fractions were subjected to Immunoblotting with indicated antibodies. **(C)** Diagram of full length Ap80 and Ap130 and FLAG-Ap80 truncations used. CC/BAR, coiled-coil/BAR domain; ABD, angiostatin binding domain; PDZ-BD, PDZ-binding domain.

### Ap80 interacts with KIF13B stalk and tail region and promotes KIF13B-PALS1 association

To map the region of KIF13B that binds Ap80-NB, we generated plasmids encoding GFP-KIF13B truncations covering most of the C-terminal stalk and tail region (Figure 2A; (Schou et al., 2017)). Co-expression with FLAG-Ap80-NB followed by FLAG IP identified residues 561-1500 (GFP-Tail 11) as the minimal Ap80-NB binding site in KIF13B (Figure 2B; Figure S1A, B); reciprocal GFP IP confirmed this result (Figure S1C). Full length FLAG-Ap80 interacted with an even shorter KIF13B fusion, GFP-Tail 8, which contains the MBS domain but lacks residues 1327-1500 present in GFP-Tail 11 (Figure 2A,C; Figure S1D), suggesting that the MBS domain is necessary for interaction with Ap80-NB and that residues 1327-1500 are also important for binding. However, the latter region is dispensable for interaction with full length Ap80. The results are consistent with our mass spectrometry data as the sequences of Tail 8 and Tail 11 are comprised within the BirAGFP-KIF13B Tail 2 fusion that interacts with AMOT (Table 1; (Serra-Marques et al., 2020)). Additionally, immunofluorescence microscopy (IFM) of RPE1 cells co-expressing FLAG-Ap80-NB or FLAG-Ap80 with different GFP-KIF13B truncations showed that those binding to FLAG-Ap80-NB (e.g. Tail 7 and 11) or FLAG-Ap80 (Tail 8) in IP (Fig. 2B, C), also co-localize with these fusions, whereas non-binding truncations (e.g. Tail 9 and 12) do not (Fig. S1E, F). Co-expression of FLAG-Ap80 with Ap80-binding GFP-KIF13B truncations increased the total cellular number of FLAG-Ap80 positive vesicles (Fig. S1G), possibly due to their fission.

**Figure 2.**
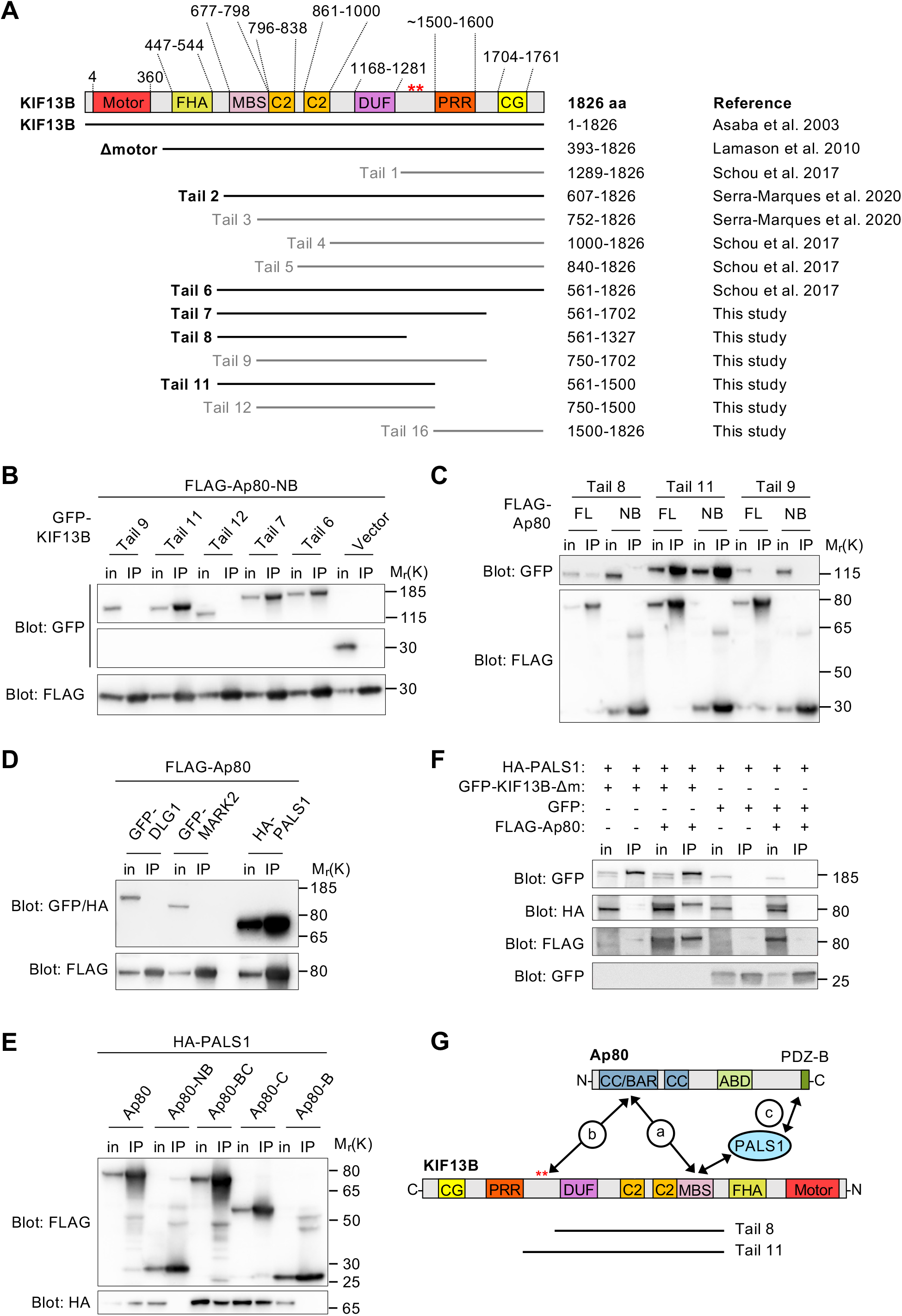
Ap80 binds KIF13B tail region and promotes KIF13B-PALS1 association. **(A)** Diagram of full length and truncated KIF13B fusions; those binding to Ap80 are in bold. FHA, Forkhead-associated domain; DUF, domain of unknown function; red asterisks, MARK2 phosphorylation sites; PRR, proline-rich region; CG, CAP-Gly domain. Redrawn from (Yoshimura et al., 2010, Soppina et al., 2014, Yamada et al., 2014, Zhu et al., 2016, Schou et al., 2017). **(B)** FLAG IP of cells co-expressing FLAG-Ap80-NB and GFP (Vector) or different GFP-KIF13B truncations. Input (in) and IP pellet (IP) fractions were analyzed by immunoblotting using indicated antibodies. **(C-F)** FLAG (**C-E**) or GFP (**F**) IP of cells expressing indicated fusion proteins, analyzed with indicated antibodies. GFP-KIF13BΔm: GFP-KIF13B-Δmotor. (**G**) Model for Ap80-KIF13B interaction.

The KIF13B MBS domain binds directly to the guanylate kinase (GUK) domains of DLG1/DLG4 and PALS1 (Yamada et al., 2007, Zhu et al., 2016), while residues 1327-1500 bind MARK2 (Yoshimura et al., 2010). We therefore asked if Ap80 binds KIF13B indirectly via these proteins. However, GFP-DLG1 and GFP-MARK2 did not co-IP with FLAG-Ap80, whereas HA-PALS1 did (Figure 2D; Figure S2A). PALS1 binds to the Ap80 C-terminus, presumably via its PDZ domain (Figure 2E; Figure S2A), while KIF13B binds to the Ap80 N-terminus (Ap80-NB; Figure 1C, D). Thus, Ap80 interacts independently with both KIF13B and PALS1. Interestingly, HA-PALS1 did not co-IP with motorless KIF13B (GFP-KIF13B-Δmotor) alone, but did upon simultaneous co-expression with FLAG-Ap80 (Figure 2F). Similar results were obtained with full-length GFP-KIF13B (Figure S2B). The inability of PALS1 to bind full-length or motorless KIF13B could be due to intramolecular interaction in KIF13B blocking access to the MBS domain. Supportively, IP using GFP- or HA-KIF13B truncations revealed that KIF13B residues 1289-1499 interact with the MBS domain (Figure S2C, D), consistent with previous work (Yamada et al., 2007). Our results suggest that Ap80 promotes association of PALS1 with KIF13B by inducing a conformational change in KIF13B that allows PALS1 to access the MBS region.

Our protein interaction data is summarized in Figure 2G. First, we suggest that the interaction (a) between the NB region of Ap80 and MBS domain in KIF13B is necessary, but not sufficient for these proteins to interact. Two additional supportive interactions can occur: one (b) between the Ap80 NB region and KIF13B residues 1327 to 1500, which seem important for a strong interaction between the proteins, and another (c) between the MBS domain and the C-terminus of Ap80, likely mediated by PALS1. Binding of Ap80 to KIF13B presumably induces a conformational change in KIF13B that permits binding of the MBS domain to PALS1.

### Co-localisation of Ap80, PALS1 and KIF13B at the base of primary cilia

Next, we asked if Ap80, KIF13B and PALS1 co-localize in cells. First, we examined the level of endogenous Ap80 and Ap130 in mouse fibroblasts (NIH 3T3 cells) and different mouse or human epithelial cell lines (IMCD3, HEK293T, RPE1 cells), by immunoblotting with AMOT antibody. We chose RPE1 cells for further analysis as they specifically expressed Ap80, but not Ap130. Further, in these cells Ap80 and KIF13B appeared to be upregulated by high cell confluency and serum deprivation (Figure S3A), conditions that promote primary cilia formation (Pugacheva et al., 2007). Previously, we showed that GFP-KIF13B localizes to and moves within primary cilia of RPE1 cells (Schou et al., 2017, Juhl et al., 2021). We examined the subcellular localization of Ap80-GFP by live imaging in serum-deprived, ciliated RPE1 cells stably expressing the ciliary membrane marker SMO-tRFP (Lu et al., 2015). We found that GFP-tagged Ap80 is concentrated at the cilia base in most cells (89% ± 8%, n=67 cells) and exhibits little, if any, movement (Figure 3A, Figure S3G, Movie 1), even when co-expressed with mCherry-KIF13B that co-localized with Ap80-GFP at the centrosome (Movie 2, Movie 3). However, we cannot rule out that the GFP tag affects Ap80 motility. In fixed cells, FLAG-Ap80 and FLAG-Ap80-NB were also detected at the ciliary base (Figure S3B, E), but recruitment of Ap80 to this site was independent of KIF13B, as centrosomal levels of FLAG-Ap80 in *KIF13B^-/-^* cells was analogous to that of wild type (WT) cells (Figure S3C, D). GFP-tagged (Figure 3B) and endogenous PALS1 (Figure S3F) were mostly dispersed localized in RPE1 cells, but upon co-expression with FLAG-Ap80 (Figure 3C) or GFP-Ap80 (Figure S3G) PALS1 accumulated at the ciliary base and at cytoplasmic Ap80-positive puncta. Furthermore, in agreement with our biochemical data GFP-Ap80 and endogenous PALS1 both co-localized with KIF13B-Δmotor-HA in intracellular puncta (Figure S3H). Together, these results indicate that Ap80 localizes to the base of cilia and where it recruits PALS1, and Ap80 is likely not a cargo of KIF13B, but Ap80, KIF13B and PALS1 co-localize within cells. In MDCK-II cells expressing PALS1 fused to APEX2-EGFP (PALS1-A2E) (Tan et al., 2020), PALS1-A2E localized around the base (Figure S3I) and proximal region of the primary cilium (Figure S3J), suggesting that such localization is conserved in cell types other than RPE1.

**Figure 3.**
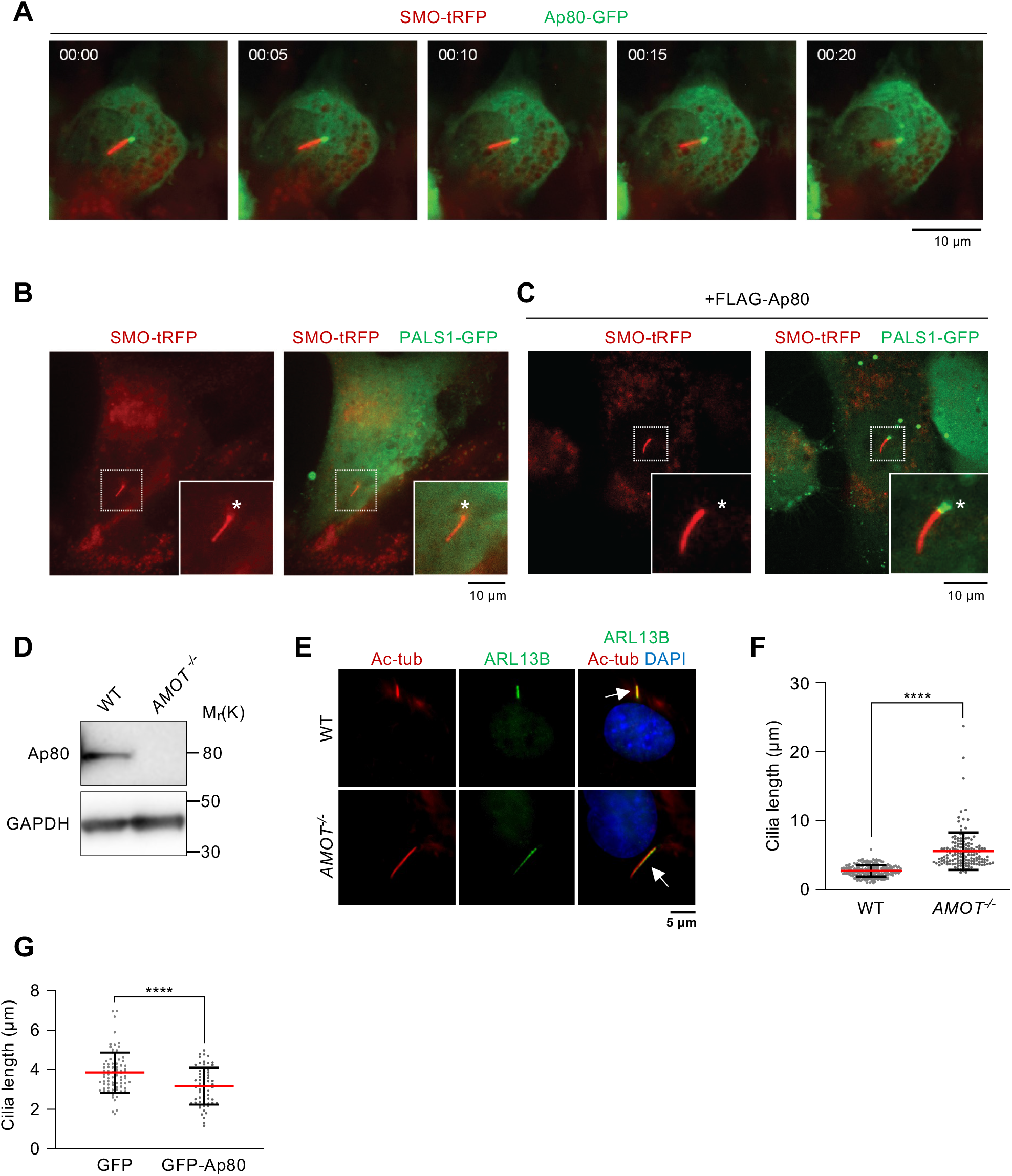
Ap80 localizes to the ciliary base and regulates ciliary length in RPE1 cells. **(A)** Frames of time-lapse movie of a cell co-expressing SMO-tRFP and Ap80-GFP. Asterisk: ciliary base. Time in seconds. (**B**, **C**) Snapshots of live cells co-expressing SMO-tRFP and PALS1-GFP in the absence (**B**) or presence (**C**) of FLAG-Ap80. Asterisk: ciliary base. **(D)** Immunoblot of WT and *AMOT^-/-^* cells using indicated antibodies. **(E)** IFM of serum-deprived WT (parental) and *AMOT^-/-^* cells using acetylated α-tubulin (AcTub) and ARL13B as cilia markers (arrows). DNA was stained with DAPI. **(F)** Cilia length quantification in WT and *AMOT^-/-^* cells; >69 cilia measured in total per condition (n=3). **(G)** Cilia length quantification in GFP or GFP-Ap80 expressing cells; 18-35 cilia measured per condition (n=3). Data in (F) and (G) show mean□±□SD. ****, P≤0.0001 (unpaired, two-tailed Student’s *t* test).

### Ap80 regulates ciliary length in RPE1 cells

Next, we set out to knockout the *AMOT* gene (coding for both Ap80 and Ap130) in RPE1 cells using CRISPR/Cas9 methodology (Ran et al., 2013). We generated a clonal *AMOT^-/-^* line containing a homozygous 1 bp deletion in the first exon resulting in a premature stop codon at position 184-186 (Figure S4A) and loss of detectable Ap80 protein (Figure 3D). Interestingly, IFM with antibodies for ciliary markers revealed that *AMOT^-/-^* RPE1 cells have significantly longer cilia than WT cells (Figure 3E, F). The *AMOT^-/-^* cells also seemed to have fewer cilia than controls, but this difference was statistically insignificant (Figure S4B). For technical reasons, rescue experiments in the *AMOT^-/-^* cells were unsuccessful, but partial depletion of Ap80 using endoribonuclease-prepared small interfering RNA (esiRNA) recapitulated the long cilia phenotype of *AMOT^-/-^* RPE1 cells (Figure S4C-E). Conversely, cells expressing GFP-Ap80 had significantly shorter cilia, but similar ciliation frequency, compared to GFP expressing cells (Figure 3G; Figure S4F). This was not simply due to protein over expression because the average relative expression level of GFP-Ap80 was about 5 times lower (18.2% ±2.0%, n=3) than that of GFP (100%, n=3), whereas average transfection efficiencies were 25.1% ±7.2% (n=3) for GFP-Ap80 and 58.8% ±6.0% (n=3) for GFP, respectively. Collectively, these results indicate that Ap80 negatively regulates ciliary length, but the underlying mechanism and functional consequences thereof remain to be clarified.

## Conclusion

We characterized Ap80 as a KIF13B interactor that promotes binding of KIF13B to PALS1. Ap80 does not appear to be a KIF13B cargo, but is localized to the ciliary base where it recruits PALS1 and controls ciliary length. Ap80 was previously shown to bind RICH1, a GTPase-activating protein for the Rho GTPases Cdc42 and Rac1 (Richnau and Aspenstrom, 2001), and mediate its targeting to a protein complex at epithelial tight junctions containing PALS1, PATJ and PAR3 (Wells et al., 2006). This promoted re-localization of PALS1, PATJ and PAR3 from tight junctions to recycling endosomes, causing loss of tight junction integrity and induction of epithelial-mesenchymal transition (Wells et al., 2006, Heller et al., 2010). Similarly, in endothelial cells Ap80 promoted breakdown of cellular junctions to stimulate endothelial to mesenchymal transition (Bratt et al., 2005, Levchenko et al., 2004, Troyanovsky et al., 2001). Our work suggests that Ap80 may have an additional role in controlling the trafficking of PALS1 or even the entire Crumbs complex (CRB3, PALS1, PATJ, LIN7C) to the ciliary base. Supportively, components of the Crumbs complex have been detected in cilia, and CRB3 is required for ciliogenesis (Fan et al., 2004, Bazellieres et al., 2018).

Although more work is needed to determine how Ap80 regulates ciliary length, and the functional consequences thereof, the mechanisms could include local regulation of actin dynamics, e.g. through ciliary recruitment of RICH1 (Wells et al., 2006). The RICH1 effector Cdc42 localizes at the ciliary base and regulates ciliary length and signaling by modulating actin dynamics, and by recruiting PAR3/PAR6/aPKC (Drummond et al., 2018). Ap80 also associates physically with the PAR3/PAR6/aPKC complex (Wells et al., 2006), which regulates ciliogenesis (Sfakianos et al., 2007, Pruliere et al., 2011) and KIF13B activity (Yoshimura et al., 2010). In future studies it will be interesting to explore these interactions further from a ciliary perspective.

## Materials and Methods

### Antibodies

For immunoblotting, the following primary antibodies were used (catalog numbers and dilutions in parenthesis): rabbit anti-FLAG (Sigma, cat# F7425; 1:1,000), rabbit anti-GFP (Santa Cruz, cat# sc-8334; 1:1,000), rabbit anti-HA (SantaCruz, cat# sc-805; 1:500), mouse anti-DCTN1 (BD Biosciences, cat# 610474; 1:1,000), mouse anti-PALS1 (Santa Cruz, cat#sc-365411; 1:200), rabbit anti-GAPDH (Cell Signaling Technology, cat# 2118S; 1:1,000), rabbit anti-AMOT (kindly provided by Dr. Joseph Kissil, Scripps Research Institute, Jupiter, FL, USA; 1:1,000), rabbit anti-IFT88 (ProteinTech, cat# 13967-1-AP; 1:2,000). Secondary antibodies used for immunoblotting were Horseradish Peroxidase (HRP)-conjugated swine anti-rabbit (DAKO, cat# P0399; 1:4,000) or goat anti-mouse (DAKO, cat# P0447; 1:4,000).

For IFM analysis, the following primary antibodies were used: mouse anti-FLAG (Sigma cat# F1804; 1:1,000), mouse anti-PALS-1 (Santa Cruz, cat#sc-365411; 1:50), goat anti-MYC (Abcam, cat# ab9132; 1:500), chicken anti-GFP (Abcam, cat# ab13970; 1:2,000), rabbit anti-ARL13B (ProteinTech cat# 17711-1-AP; 1:500), rabbit anti-AMOT (kindly provided by Joseph Kissil, Scripps Research Institute, Jupiter, FL, USA; 1:200), mouse anti-DCTN1 (BD Biosciences, cat# 610473; 1:500), mouse anti-acetylated α-tubulin (Sigma, cat# T7451; 1:2,000), rabbit anti-CEP164 (Sigma, cat# HPA037606; 1:500). Secondary antibodies used for IFM were all from Invitrogen and diluted 1:600: Alexa Fluor 488-conjugated donkey anti-mouse (A-21202), donkey anti-goat (A-11055), donkey anti-rabbit (A-21206) and goat anti-chicken (A11039); Alexa Fluor 568-conjugated donkey anti-mouse (A-10037), and donkey anti-rabbit (A-10042); Alexa Fluor 647-conjugated donkey anti-mouse (A-31571), donkey anti-rabbit (A-31573).

### PCR, cloning procedures and plasmids

All PCR and cloning was done following standard procedures; primer sequences are listed in Table S1 and plasmids used are listed in Table S2. Plasmids encoding truncated versions of GFP-KIF13B (Tail 7, 8, 9, 11 and 12) were generated by PCR with relevant primers using pEGFP-C1-KIF13B as template (Asaba et al., 2003), followed by cloning into pEGFP-C1 (Clontech). Plasmids encoding truncated KIF13B-HA fusions (Tail 1 and 16) were generated by PCR with relevant primers using pcDNA3-KIF13B-HA plasmid (Lamason et al., 2010) as template. Plasmids encoding truncated versions of FLAG-Ap80 (-NB, -B, -C and -BC) were generated by PCR with relevant primers using pCMV-Ap80 as template, followed by cloning into pFLAG-CMV2 (Sigma). Plasmid encoding GFP-Ap80 or Ap80-GFP was generated by PCR with relevant primers using pCMV-Ap80 as template, followed by cloning into pEGFP-C1 or pEGFP-N1, respectively (Clontech). For CRISPR-Cas9 mediated knockout of AMOT, 25 nt oligos targeting exon 1 in AMOT (5’-AATACCGTGGTCCCTCCACTTGG-3’) were cloned into pSpCas9(BB)-2A-Puro (PX459) using the procedure in (Ran et al., 2013). *Escherichia coli* DH10B was used for transformation and plasmid amplification. Plasmid purification was performed using NucleoBond Xtra Midi EF Kit from Macherey-Nagel. Inserts were sequenced at Eurofins MWG Operon, Ebersberg, Germany.

### Cell culture and transfections

Unless otherwise stated, cells were grown at 37°C, 5% CO_2_ and 95% humidity in Dulbecco’s modified Eagle’s medium (DMEM, Gibco) with 10% heat inactivated fetal bovine serum (FBS, Gibco) and 10 ml 1^-1^ penicillin-streptomycin (Gibco) and cell cultures were passaged every 3-4 days. Cell lines used were: HEK293T cells (laboratory stock, originally derived from American Type culture collection (ATCC), clone CRL-3216); hTERT-RPE1 cells (laboratory stock; originally derived from the immortalized hTERT-RPE1 cell line, ATCC, clone CRL-4000); hTERT-RPE1 cells lacking KIF13B (*KIF13B*^-/-^; (Schou et al., 2017)); hTERT-RPE1 cells stably expressing SMO-tRFP (Lu et al., 2015); swiss NIH3T3 mouse fibroblasts (laboratory stock, originally derived from ATCC clone CRL-1658); Mouse Inner Medullary Collecting Duct (IMCD3) cells (laboratory stock, originally derived from ATCC, clone CRL-2123); and PALS1-A2E MDCK-II cells (Tan et al., 2020). IMCD3 cells were grown in 45% DMEM and 45% F-12 (Ham; Sigma) with 10% FBS and 10 ml 1^-1^ penicillin-streptomycin whereas the PALS1-A2E MDCK-II cells were grown and subjected to APEX2-mediated biotinylation as described previously (Tan et al., 2020). Generation of *AMOT*^-/-^ RPE1 clone was done using the CRISPR-Cas9 method described in (Ran et al., 2013) with plasmid pSpCas9(BB)-2A-Puro (PX459) encoding AMOT-specific guide RNA targeting the first exon of AMOT (5’-AATACCGTGGTCCCTCCACTTGG-3’). Screening of 91 clones by immunoblot analysis led to identification of a single clone with no detectable Ap80 expression. Sequencing of PCR-amplified genomic DNA from this clone revealed a 1 bp deletion in exon 1 resulting in a premature stop codon. All cell lines were routinely tested for contamination.

For plasmid transfections cells were grown to 40-50% confluency and transfections were performed using FuGene6 (Promega, E2692) according to the manufacturer’s protocol. For IP experiments, HEK293T cells were grown in 10 cm diameter petri dishes and transfected with 4 μg DNA. Six hours after transfection the medium was changed to fresh growth medium and the cells were incubated for 16 hrs before harvest. For IFM experiments, RPE1 cells were grown on glass coverslips in 20 mm diameter dishes and transfected with 1 μg DNA. Six hours after transfection the medium was changed to either growth or serum-deprived medium for 16-24 hrs. For immunoblot experiments, RPE1 cells were grown in 60 mm diameter dishes and transfected with 2 μg DNA. Six hours after transfection the medium was changed to either growth or serum-deprived medium for 16-24 hrs.

All transfections with esiRNA (Sigma MISSION^®^ esiRNA human AMOT cat# EHU129771) and control siRNA (5’-UAAUGUAUUGGAAUGCAUA(dTdT)-3’ from Eurofins MWG Operon) were carried out as double-transfections using DharmaFECT Duo (Dharmacon). RPE1 cells were grown to 80% confluency and transfected according to the manufacturer’s manual. Six hours after transfection the medium was changed to fresh growth medium. The following day the cells were transfected again. 36 hrs later, the cells were split and an appropriate number of cells were seeded for IFM or immunoblot analysis. Prior to fixation or cell lysis the cells were serum-deprived for 16-24 hours.

### Streptavidin pulldown assays and mass spectrometry

HEK293 cells were transfected with BirAGFP-KIF13B Tail 2 or Tail 3 constructs (designated C2 and C3, respectively, in (Serra-Marques et al., 2020)) using Polyethylenimine (PEI; Mw 2500; Polysciences) at a 3:1 PEI:DNA ratio (w/w). Cells were harvested 24 hours after transfection, by scraping the cells in ice-cold PBS and lysing cell pellets in the lysis buffer (20 mM Tris-HCl, pH 7.5, 100 mM NaCl, 1.0% Triton X-100, and protease inhibitors; Roche). Supernatants and pellet fractions were separated by centrifugation at maximum speed for 20 minutes. Supernatants were mixed with an equal amount of Dyna M-280 Streptavidin beads (Life Technologies). Samples were incubated for 2 hours while rotating at 4°C, collected with a magnet and pellets were washed 5-7 times with the wash buffer (20 mM Tris-HCl, pH 7.5, 100 mM NaCl, 0.1 % Triton X-100). Samples were eluted in SDS sample buffer and 30 μl of each sample was run on a 12% Bis-Tris 1D SDS-PAGE gel (Biorad) for 1 cm and stained with colloidal Coomassie dye G-250 (Gel Code Blue Stain Reagent, Thermo Scientific). Each lane was cut into 1 band, which was treated with 6.5 mM dithiothreitol (DTT) for 1 hour at 60 °C for reduction and 54 mM iodoacetamide for 30 min for alkylation. The proteins were digested overnight with trypsin (Promega) at 37°C. The peptides were extracted with acetonitrile (ACN) and dried in a vacuum concentrator. The data were acquired using an Orbitrap Q Exactive mass spectrometer. Peptides were first trapped (Dr Maisch Reprosil C18, 3 μm, 2 cm x 100 μm) before being separated on an analytical column (Zorbax SB-C18, 1.8 μm, 40 cm x 50 μm), using a gradient of 60 min at a column flow of 150 nl min^-1^. Trapping was performed at 8 μl/min for 10 min in solvent A (0.1 M acetic acid in water) and the gradient was as follows 7-30% solvent B (0.1 M acetic acid in acetonitrile) in 31 min, 30-100% in 3 min, 100% solvent B for 5 min, and 7% solvent B for 13 min. Full scan MS spectra from m/z 350-1500 were acquired at a resolution of 35,000 at m/z 400 after accumulation to a target value of 3e6. Up to ten most intense precursor ions were selected for fragmentation. HCD fragmentation was performed at normalized collision energy of 25% after the accumulation to a target value of 5e4. MS/MS was acquired at a resolution of 17,500. In all cases nano-electrospray was performed at 1.7 kV using an in-house made gold-coated fused silica capillary (o.d. 360 μm; i.d. 20 μm; tip i.d. 10 μm). Raw files were processed using Proteome Discoverer 1.3 (Thermo Scientific, Bremen, Germany). The database search was performed against the Swissprot human database, taxonomy (version November 2012) using Mascot (version 2.3, Matrix Science, UK) as search engine. Carbamidomethylation of cysteines was set as a fixed modification and oxidation of methionine was set as a variable modification. Trypsin was specified as enzyme and up to two miss cleavages were allowed. Data filtering was performed using percolator, resulting in 1% false discovery rate (FDR). Additional filter was Mascot ion score >20. Raw files corresponding to one sample were merged into one result file.

### Immunoprecipitation

24 hours before IP HEK293T cells were transfected with relevant plasmids. Cells were harvested using ice cold EBC buffer (140 mM NaCl, 50 mM Tris-HCl, 0.5% NP-40, 5 mM EDTA and protease inhibitor cocktail (Roche)) and briefly sonicated prior to 20 min centrifugation at 20,000 *g*, 4°C. For FLAG, GFP and HA IP, cleared cell lysates were incubated rotating for 1 hr at 4°C with 20 μl bead slurry of Anti-FLAG (M2) conjugated beads (Sigma, cat# A2220) or 25 μl bead slurry of GFP-Trap^®^_A beads (Chromotek, cat# gta-20) or 20 μl bead slurry of Pierce^™^ Anti-HA Agarose Conjugate (Thermo Fisher Scientific, cat#26181), respectively. The immunocomplexes were washed 4 times with EBC buffer and eluted with SDS-PAGE sample buffer. The eluted proteins were then subjected to SDS-PAGE and immunoblotting. All IP experiments were repeated independently at least three times, and figures show representative results from these independent repeats.

### Protein concentration determination, SDS-PAGE and immunoblotting

The BioRad *DC* Protein Assay was used to determine the protein concentration of cell lysates. SDS-PAGE and immunoblotting was performed as described in (Schou et al., 2017).

### Immunofluorescence microscopy

For IFM analysis, RPE1 cells were grown on glass coverslips, and subjected to 16-24 hrs serum-deprivation to induce ciliogenesis. Cells were washed in ice cold PBS, fixed with 4% paraformaldehyde (PFA) solution for 15 min at room temperature (RT) and cell membranes were permeabilized by incubation in 1x PBS with 0.2% (v/v) Triton X-100 and 1% (w/v) bovine serum albumin (BSA) for 12 min at RT. A blocking step of 30 min incubation at RT in 1x PBS with 2% BSA was done to avoid unspecific binding of antibodies, followed by incubation with primary antibodies overnight at 4°C. After three washing steps with BSA, cells were incubated in appropriate Alexa Fluor-conjugated secondary antibodies, diluted in BSA, for 45 min at RT and nuclei were labeled with DAPI. Coverslips were mounted using 1x PBS, 90% glycerol and 2% n-Propyl Gallate on glass slides and edges were sealed with nail polish. Fluorescence images were captured on a fully motorized Olympus BX63 upright microscope with an Olympus DP72 color, 12.8-megapixel, 4.140 x 3.096-resolution camera and with a fully motorized and automated Olympus IX83 Inverted microscope with a Hamamatsu ORCA-Flash 4.0 camera (C11440-22CU). The software used was Olympus CellSens dimension. Images were processed for publication using Adobe Photoshop version CS6. For quantifications of the centrosomal FLAG-Ap80 levels images were analyzed using ImageJ software. An outline was drawn around each centrosome and using the measurement- and integrated density (IntDens) functions, the Mean Green Fluorescence Intensity was measured in this area along with a background reading. The corrected mean fluorescence intensity in the centrosome was calculated by subtracting the corresponding background value. Procedures for confocal microscopy analysis of PALS1-A2E MDCK-II cells are described in (Tan et al., 2020).

### Live cell imaging

Transfection and live cell imaging of RPE1 cells stably expressing SMO-tRFP were done as described previously (Juhl et al., 2021). Live cell imaging of Ap80-GFP and mCherry-KIF13B was performed using Total Internal Reflection Fluorescence (TIRF) microscopy. Inverted research microscope Nikon Eclipse Ti-E (Nikon), equipped with the perfect focus system (Nikon), Nikon Apo TIRF 100x N.A. 1.49 oil objective (Nikon) and iLas3 system (Dual Laser illuminator for azimuthal spinning TIRF (or Oblique) illumination and Simultaneous Targeted Laser Action including PhotoAblation; Gataca Systems) was used. The system was also equipped with ASI motorized stage MS-2000-XY (ASI), Photometrics Evolve 512 Delta EMCCD back illuminated camera (Teledyne Photometrics) and controlled by the MetaMorph 7.8 software (Molecular Devices). Stradus 488 nm (150 mW, Vortran) and OBIS 561 nm (100 mW, Coherent) lasers were used as the light sources. We used ZT405/488/561/640rpc ZET405/488/561/635m filter set (TRF89901, Chroma) together with Optosplit III beamsplitter (Cairn Research Ltd, UK) equipped with double emission filter cube configured with ET525/50m, ET630/75m and T585LPXR (Chroma). 16-bit images were projected onto the EMCCD chip with intermediate lens 2.5X (Nikon C mount adapter 2.5X) at a magnification of 0.065 μm/pixel. To keep cells at 37°C we used stage top incubator (model INUBG2E-ZILCS, Tokai Hit).

### Statistical analysis

Statistical analysis was performed using GraphPad Prism 6 software.

## Acknowledgements

We thank Søren Johansen, Emilie Baldram, Ida Hemmingsen, Signe Aagard and Sara Enevoldsen for technical assistance, and Joseph Kissil, Athar Chisthi, Joel Pomerantz, Sadanori Watanabe, Hiroaki Miki, Ronald Roepman, Francesc Garcia-Gonzalo, Lars Ellgaard, Dannel McCollum, Christopher Westlake, and Jeffrey Miner for reagents. Supported by grants from Novo Nordisk Foundation (NNF14OC0011535, NNF15OC0016886), Hartmann Fonden (A31662), Danish Cancer Society (R146-A9590-16-S2), the University of Copenhagen (UCPH) Excellence Programme for Interdisciplinary Research (LBP, STC), Department of Biology, UCPH (SKM), and the Netherlands X-omics Initiative partially funded by NWO (project 184.034.019 to AA).

