## Supplemental figures, tables and legends for "Angiomotin isoform 2 promotes binding of PALS1 to KIF13B at the base of primary cilia and suppresses ciliary elongation"

### Supplementary information

### Supplementary figures

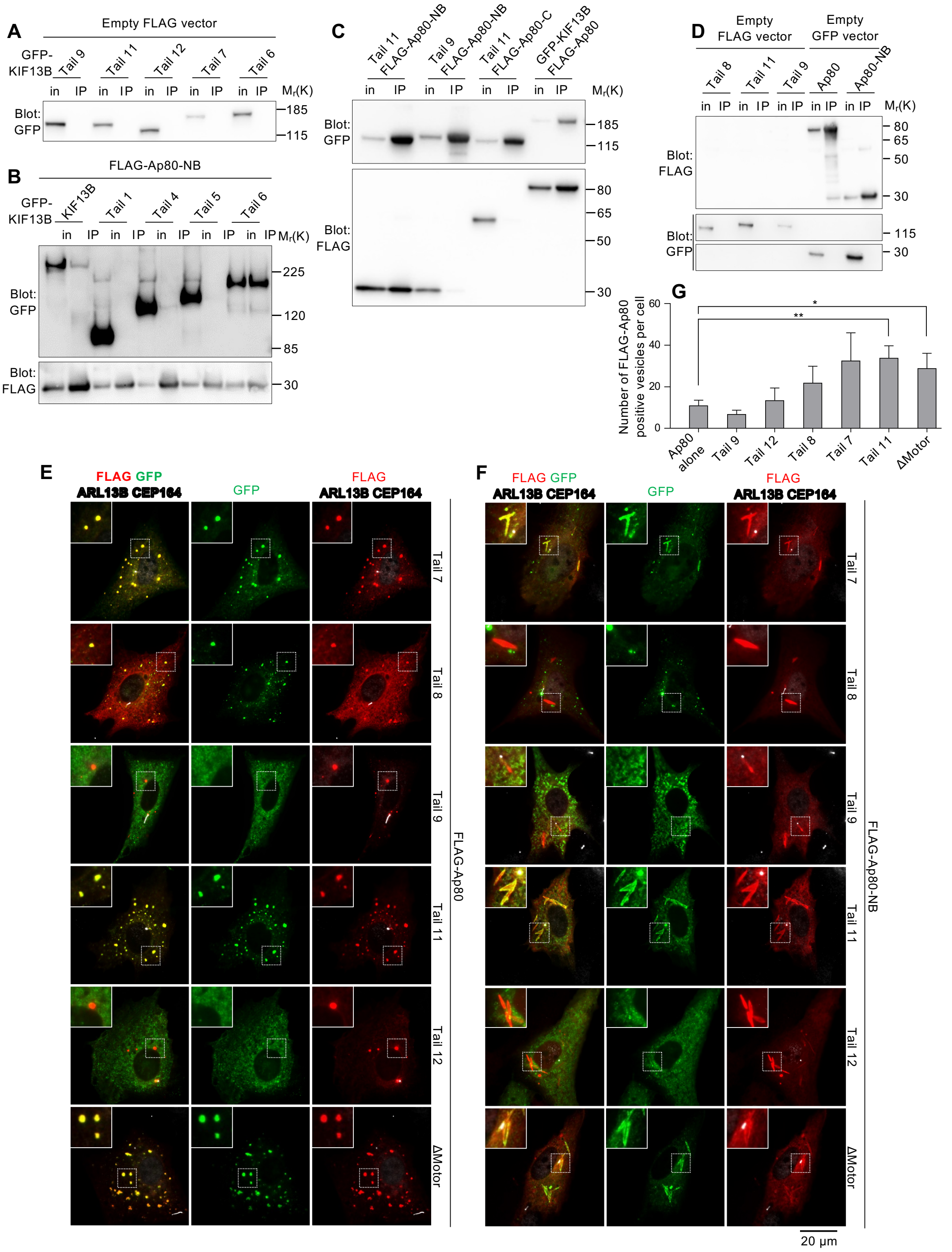


**Figure S1. Control experiments for Figure 2. (A)** FLAG IP analysis of HEK293T cells co-expressing empty FLAG vector and the indicated GFP-KIF13B tail fusions. Input (in) and IP pellets (IP) were subjected to SDS-PAGE and immunoblot analysis using GFP antibody. **(B)**

FLAG IP analysis as in (B) using HEK293T cells co-expressing FLAG-Ap80-NB and either full length GFP-KIF13B (KIF13B) or the indicated GFP-KIF13B tail fusions. **(C)** GFP IP analysis of HEK293T cells co-expressing different combinations of FLAG-Ap80 fusions and GFP-KIF13B tail fusions as indicated. GFP fusions were IPed using anti-GFP conjugated beads and input (in) and IP pellets (IP) subjected to immunoblot analysis using indicated antibody. Note that the two last lanes are the same as those depicted in Figure 1B. **(D)** FLAG IP analysis of HEK293T cells co-expressing empty FLAG vector with indicated GFP-KIF13B tail fusions and cells co-expressing GFP alone with either FLAG-Ap80 or FLAG-Ap80-NB. FLAG fusions were IPed using anti-FLAG conjugated beads and input (in) and pelleted proteins (IP) subjected to immunoblot analysis using indicated antibody. **(E, F)** IFM analysis of RPE1 cells co-expressing FLAG-Ap80 **(E)** or FLAG-Ap80-NB **(F)** and different GFP-KIF13B tail fusions as indicated. Antibodies against ARL13B and CEP164 (pseudocolored white) mark the cilium-centrosome axis. Insets show enlargements of regions with vesicles or tubular structures induced by FLAG-Ap80/Ap80-NB. **(G)** Quantification of the number of FLAG-Ap80 positive vesicles per cell expressing FLAG-Ap80 alone or together with different GFP-KIF13B tail fusions (21-50 cells analyzed per condition, n=3, except for Δmotor where n=2). Bars represent mean ±SD. * and **; $P \leq 0.05$ and 0.01, respectively (one-way ANOVA).

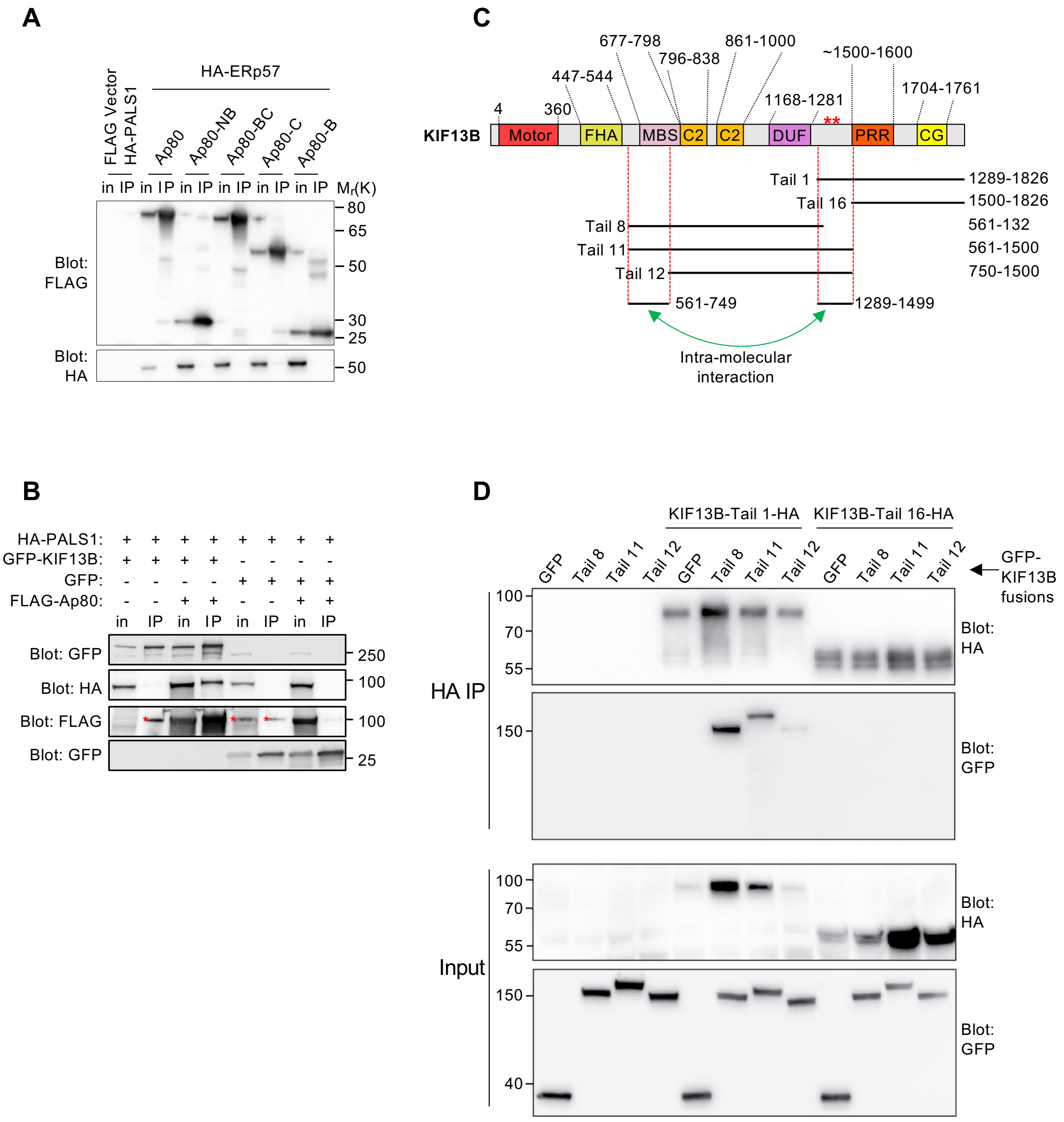


**Figure S2. Control IP experiments for Figure 2. (A)** Control experiment for Figure 2E. FLAG IP analysis of HEK293T cells co-expressing HA-ERp57 and the indicated FLAG-Ap80 constructs and cells co-expressing empty FLAG vector and HA-PALS1. **(B)** GFP IP analysis of HEK293T cells expressing the indicated fusion proteins. Input (in) and IP pellet (IP) fractions were analyzed by immunoblotting using indicated antibodies. **(C, D)** Schematics of GFP- or HA-tagged KIF13B tail fragments **(B)** used for HA IP analysis in HEK293T cells, followed by immunoblotting with GFP or HA antibodies as indicated **(D)**. Note that KIF13B-Tail 1-HA (residues 1289-1826) binds to GFP-Tail 8 and 11 that both contain the MBS domain whereas KIF13B-Tail 16-HA (residues 1500-1826) does not.

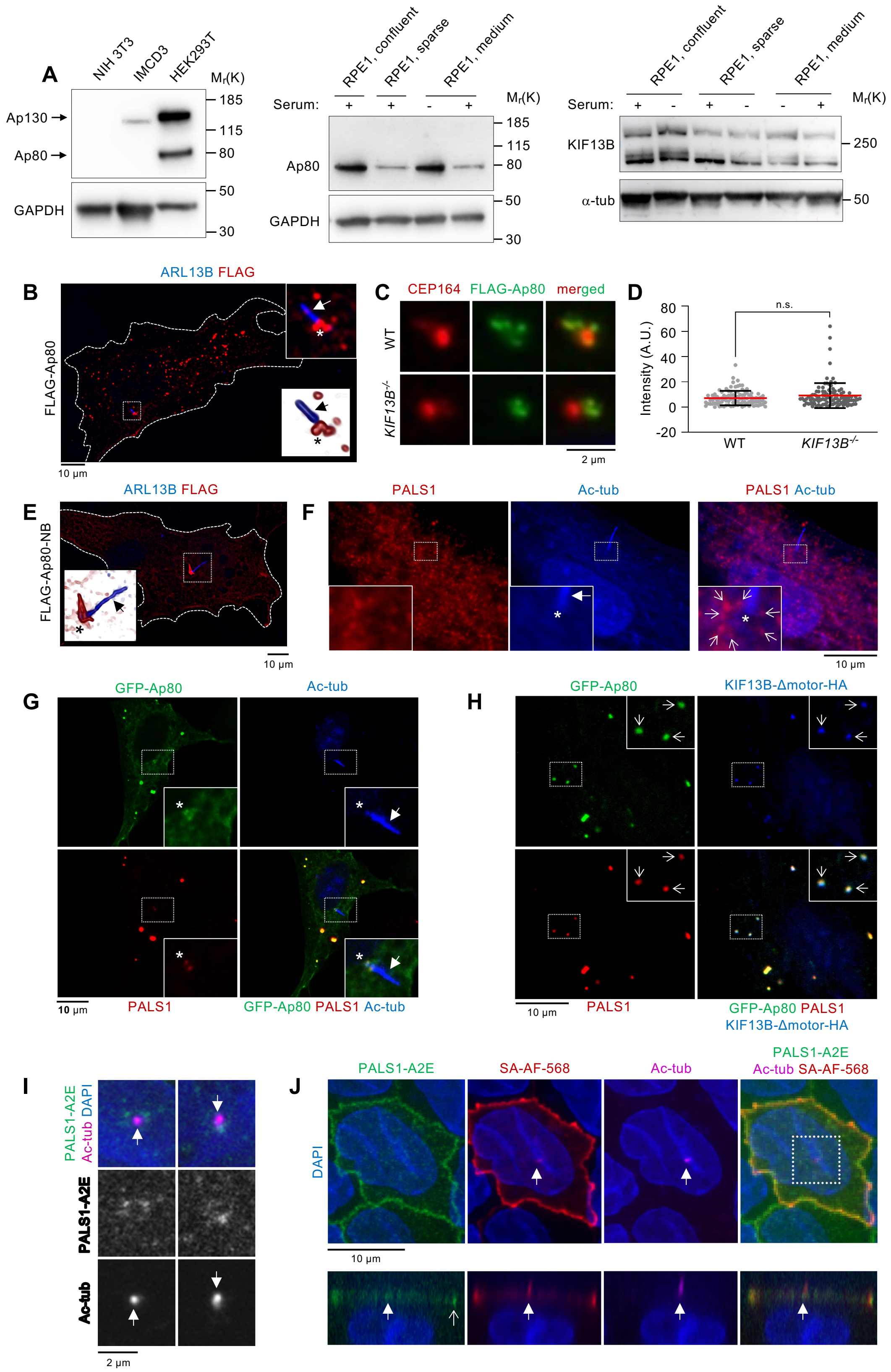


**Figure S3. Ap80, KIF13B and PALS1 localization in ciliated RPE1 cells. Related to Figure 3. (A)** Immunoblot analysis of cell lysates from NIH3T3, IMCD3, HEK293T (left panel), and

RPE1 (middle and right panel) cells using antibodies as indicated. RPE1 cells were grown with (+) or without (-) serum and to indicated confluency. KIF13B migrates as a double band in these cells as shown before (Schou et al., 2017). **(B, C)** IFM of FLAG-Ap80 in RPE1 cells using indicated antibodies. In (B) ARL13B marks the cilium (arrows), and asterisks indicate the ciliary base. In (C), transfected WT and *KIF13B*$^{-/-}$ cells (Schou et al., 2017) were analyzed in parallel and CEP164 antibody was used to mark the centrosome/mother centriole. **(D)** Quantification of relative centrosomal levels of FLAG-Ap80 in WT and *KIF13B*$^{-/-}$ cells, based on images as in (C). Between 29 and 39 cells were analyzed per condition (n=3). Bars represent mean ± SD. P value results from unpaired two-tailed t-test (n.s., $P \geq 0.05$, not significant). **(E)** IFM of FLAG-Ap80-NB in RPE1 cells using indicated antibodies; ARL13B marks the cilium (arrows). Asterisks indicate the ciliary base. Inset in (C) shows 3D isosurface rendering on captured z stacks of the boxed area. **(F)** IFM of untransfected cells using antibodies against acetylated α-tubulin (Ac-tub) and PALS1. Closed arrow and asterisk mark the cilium and centrosome, respectively, while open arrows indicate periciliary staining of PALS1 in the enlarged insert. The image shown is representative of 39 ciliated cells examined. **(G, H)** IFM of cells expressing GFP-Ap80 without **(G)** and with co-expression of KIF13B-ΔMotor-HA **(H)** using antibodies as indicated. Closed arrows point to cilia; asterisks mark the ciliary base; open arrows point to cellular puncta where GFP-Ap80, KIF13B-ΔMotor-HA and endogenous PALS1 co-localize. In (G, H), 74% of the GFP-Ap80 puncta (n=80) were positive for endogenous PALS1; in (H) 53% of GFP-Ap80 puncta were positive for both PALS1 and KIF13B-ΔMotor-HA, and 21% were positive for KIF13B-ΔMotor-HA but not PALS1 (n=57). We did not observe co-localization of KIF13B-ΔMotor-HA and PALS1 in the absence of GFP-Ap80. **(I)** Confocal images of filter-grown PALS1-A2E MDCK-II cells (Tan et al., 2020) stained with acetylated tubulin antibody (magenta) to mark the cilium (closed arrow). DNA is stained with DAPI. **(J)** Confocal images of filter-grown PALS1-A2E MDCK-II cells subjected to APEX2-mediated proximity biotinylation for 1 min, followed by staining with fluorescently labeled streptavidin (SA-AF-568; red) and acetylated tubulin antibody (magenta) to mark the cilium (closed arrow). Upper and lower panels show the same cell viewed from the top and side, respectively. Open arrow shows PALS1 staining at the cell-cell junction.

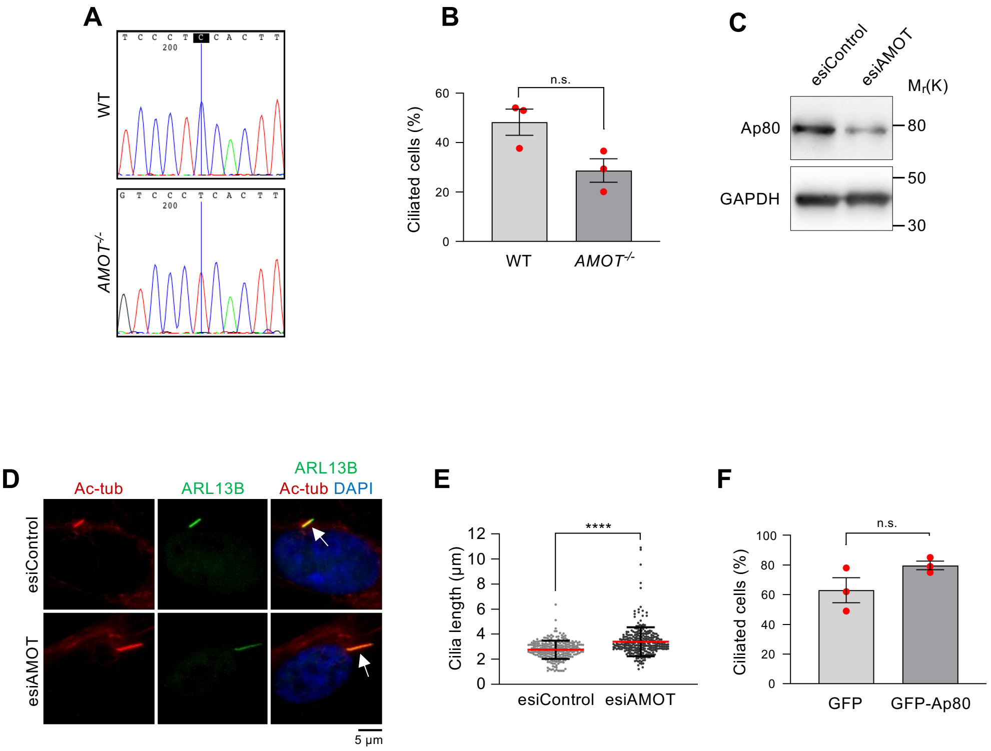


**Figure S4. Characterization of AMOT-depleted cells. Related to Figure 3. (A)** Chromatogram illustrating the genomic sequence of WT control RPE1 cells (upper panel) and the *AMOT*$^{-/-}$ clone (lower panel) in the region targeted by CRISPR/Cas9. The marked cytosine is deleted in the *AMOT*$^{-/-}$ clone, causing a premature stop codon. **(B)** Quantification of ciliation frequency of WT (parental) and *AMOT*$^{-/-}$ RPE1 cells. At least 100 cells were analyzed per condition per experiment (n=3). Data are represented as mean ± SD and significance was determined using an unpaired, two-tailed Student's *t* test (n.s., P ≥ 0.05, not significant). **(C)** Immunoblot analysis of WT RPE1 cells treated with esiAMOT or control esiRNA. Blots were probed with indicated antibodies. Quantification of band intensities showed an average Ap80 protein level of 49.1% ±11.6% in esiAMOT treated cells relative to esiControl treated cells (n=3). **(D)** IFM analysis of serum-deprived WT cells treated with esiAMOT or control esiRNA. Cells were stained with antibodies against acetylated α-tubulin (AcTub, red) and ARL13B (green) to mark the cilium (arrows). DNA was stained with DAPI (blue). **(E)** Quantification of cilia length based on data as shown in (D), with at least 69 cilia measured in total per condition (n=3). Data are represented as mean ± SD and significance was determined using an unpaired, two-tailed Student's *t* test: ****, P≤0.0001. (**F**) Quantification of ciliation frequencies in RPE1 cells expressing GFP or GFP-Ap80. Between 32-102 transfected cells were analyzed per condition per experiment (n=3). Data are represented as mean ± SD and significance was determined using an unpaired, two-tailed Student's *t* test (n.s., P ≥ 0.05, not significant).

#### Supplementary tables

##### Table S1. Primers used in this study.

| Primer name | Sequence 5'-3' * | Used for |
|---|---|---|
| Ap80F | CCGAATTCTATGCCTCGGGCTCAGCCA | FLAG-Ap80-NB<br>GFP-Ap80 |
| Ap80F_HindIII | CCAAGCTTATGCCTCGGGCTCAGCCA | Ap80-GFP |
| Ap80(735)R | CAGGATCCTTATTCTGAAACGTTGGTGGGCTG | FLAG-Ap80-NB<br>FLAG-Ap80-B |
| Ap80(106)F | CCGAATTCTAACCGGAACTTGAGGCAAGA | FLAG-Ap80-BC<br>FLAG-Ap80-B |
| Ap80FL_R | CAGGATCCTTAGATGAGATATTCCACCATCTC | FLAG-Ap80-BC<br>FLAG-Ap80-C<br>GFP-Ap80 |
| Ap80_R_BamH1 | AAGGATCCGCGATGAGATATTCCACCATCTCTGCA | Ap80-GFP |
| Ap80(670)F | ACGAATTCTAGGGAACTGGAATCCCTGAG | FLAG-AP80-C |
| 13B_561F | CCGGTACCTCCATGAAGAACGAGAATAGTTC | GFP-Tail 7, 8, 11 |
| KIF13B_1702R (13B_5106R) | AAGGATCCTCATCGGAGCCACTCCGG | GFP-Tail 7, 9 |
| 13B_3981R | CAGGATCCTCAATTGGCTGCCATTCTTGCTA | GFP-Tail 8 |
| KIF13B_750F | AAAAAGGTACCATTTGGTCTTTGGAAAAACTGGAC | GFP-Tail 9, 12 |
| KIF13B_1500R | AAGGATCCTCAGATGTCCGGGCTGGCTG | GFP-Tail 11, 12 |
| ZAN_030 | AAAAGGTACCATGTCTCATCGAAGTTCTATTCCT | Tail 1-HA |
| ZAN_102 | AAAAGGTACCATGATCAGGGTGACCAGGATGG | Tail 16-HA |
| ZAN_031 | AAAATCTAGATTATCAAGCGTAATCTGGAACATCGTATGGGTAGCTGGCCCAGGATTTC | Tail 1-HA, Tail 16-HA |
| Ap130 1r | CAGTGCTAGGGCCTGGAATC | Sequencing of *AMOT*$^{-/-}$ RPE1 cells |
| Ap130 2f | GACTGGATTTCCTGCCCCTG | Sequencing of *AMOT*$^{-/-}$ RPE1 cells |

* Relevant restriction endonuclease sites are underlined.

**Table S2. Plasmids used in this study.**

| Vector | Protein product encoded by insert* | Reference/source |
|---|---|---|
| pEGFP-C1 | none | Clontech |
| pEGFP-N1 | none | Clontech |
| pFLAG-CMV2 | none | Sigma |
| pCMV-FLAG | Ap130 | (Yi et al., 2013) |
| pCMV-FLAG | Ap80 | (Yi et al., 2013) |
| pFLAG-CMV2 | Ap80 aa 1-245 (NB) | This study |
| pFLAG-CMV2 | Ap80 aa 35-675 (BC) | This study |
| pFLAG-CMV2 | Ap80 aa 222-675 (C) | This study |
| pFLAG-CMV2 | Ap80 aa 35-245 (B) | This study |
| pEGFP-C1 | Ap80 | This study |
| pEGFP-N1 | Ap80 | This study |
| pEGFP | DLG1 (rodent) | Gift from Jeffrey Miner |
| pcDNA3 | HA-ERp57 | (Appenzeller-Herzog et al., 2008) |
| pEGFP-C1 | KIF13B | (Asaba et al., 2003) |
| Modified pEGFP-C1 vector with BirA insertion | KIF13B aa 607-1826 (Tail 2; designated C2 in (Serra-Marques et al., 2020)) | (Lansbergen et al., 2006, Serra-Marques et al., 2020) |
| Modified pEGFP-C1 vector with BirA insertion | KIF13B aa 752-1826 (Tail 3; designated C3 in (Serra-Marques et al., 2020)) | (Lansbergen et al., 2006, Serra-Marques et al., 2020) |
| pEGFP-C1 | KIF13B aa 1289-1826 (Tail 1) | (Schou et al., 2017) |
| pEGFP-C1 | KIF13B aa 1000-1826 (Tail 4) | (Schou et al., 2017) |
| pEGFP-C1 | KIF13B aa 840-1826 (Tail 5) | (Schou et al., 2017) |
| pEGFP-C1 | KIF13B aa 561-1826 (Tail 6) | (Schou et al., 2017) |
| pEGFP-C1 | KIF13B aa 561-1702 (Tail 7) | This study |
| pEGFP-C1 | KIF13B aa 561-1327 (Tail 8) | This study |
| pEGFP-C1 | KIF13B aa 750-1702 (Tail 9) | This study |
| pEGFP-C1 | KIF13B aa 561-1500 (Tail 11) | This study |

| | | |
|---|---|---|
| pEGFP-C1 | KIF13B aa 750-1500 (Tail 12) | This study |
| pcDNA3 | KIF13B-Δmotor-eGFP | (Lamason et al., 2010) |
| pcDNA3 | KIF13B-HA | (Lamason et al., 2010) |
| pcDNA3 | KIF13B-Δmotor-HA | (Lamason et al., 2010) |
| pcDNA3.1 | KIF13B aa 1289-1826 (Tail 1-HA) | This study |
| pcDNA3.1 | KIF13B aa 1500-1826 (Tail 16-HA) | This study |
| pBa-eGFP | MARK2 | Addgene #66706 |
| Gateway plasmid | 3xHA-PALS1 | (Gosens et al., 2007) |
| pEGFP-N1 | PALS1 | (Tan et al., 2020) |
| pSpCas9(BB)-2A-Puro (PX459) | 25 nt oligo targeting exon 1 in AMOT: 5'-AATACCGTGGTCCCTCCACTTGG -3' | This study; (Ran et al., 2013) |

*Unless otherwise indicated, names refer to human proteins.

#### Supplementary Movie Legends

**Movie 1. Example of live imaging of Ap80-GFP in RPE1 cells stably expressing SMO-tRFP.** Time-lapse movie of a cell co-expressing SMO-tRFP and Ap80-GFP, recorded on a spinning disk confocal microscope, with time indicated in seconds.

**Movie 2 and Movie 3. Examples of live imaging of mCherry-KIF13B and Ap80-GFP.** RPE1 cells co-expressing Ap80-GFP and mCherry-KIF13B were imaged using TIRF microscopy. No colocalization between Ap80-GFP and mCherry-KIF13B was detected except at the centrosome region. Scale bar 5 µm. 5 frames per second stream imaging with 20 ms exposure.
